# TCPGdb: A comprehensive T cell perturbation genomics database for identification of critical T cell regulators

**DOI:** 10.1101/2024.12.30.630773

**Authors:** Chuanpeng Dong, Feifei Zhang, Kaiyuan Tang, Nipun Verma, Xinxin Zhu, Di Feng, James Cai, Hongyu Zhao, Sidi Chen

**Affiliations:** Department of Genetics, Yale University School of Medicine, New Haven, CT, USA; System Biology Institute, Yale University, West Haven, CT, USA; Center for Cancer Systems Biology, Yale University, West Haven, CT, USA; Center for Biomedical Data Science, Yale University School of Medicine, New Haven, CT, USA; Yale-Boehringer Ingelheim Biomedical Data Science Fellowship Program; Yale Comprehensive Cancer Center, Yale University School of Medicine, New Haven, CT, USA; Department of Biostatistics, Yale University School of Public Health, New Haven, CT, USA; Global Computational Biology and Digital Sciences, Boehringer Ingelheim Pharmaceuticals, Inc., Ridgefield, CT, USA

## Abstract

CAR-T therapies utilizing T cells engineered with chimeric antigen receptors (CARs) have revolutionized the treatment of hematologic and immune-related malignancies. However, the anti-tumor capability of engineered CAR-T cells are often not persistent, which greatly dampen its clinical efficacy. To address this limitation, massive parallel genetic screens were widely used to identified novel targets and regulators that enhance T cell anti-tumor capability and persistence in tumor microenvironment. We hypothesized that by combining the pooled screen data from multiple independent genetic screens we could provide a systematic, comprehensive, and robust analysis of the effect of gene perturbation on T-cell based immunotherapies. After collecting data from previously published T cell screens, including CRISPR-based and ORF-based screens, through Gene Expression Omnibus (GEO), we reprocessed the gene hits summary and conducting a pathway enrichment analysis. A T cell screen perturbation score (TPS) was employed to quantifies the impact of the gene on T cell function. Additionally, gene expression data (both bulk RNA level and single-cell RNA level) from autoimmune disease and T cell-derived cancers were analyzed to pick up gene perturbations that potentially augment T cell proliferation. We integrated all data and analysis on 27 T-cells screens into our state-of-the-art T cell perturbation genomics database (TCPGdb), which is made easily accessible through our webserver and allows users to interactively explore the impact of query genes on T cell function based on prior screen data and our TPS scoring module. TCPGdb is publicly accessible at http://tcpgdb.sidichenlab.org/.

## Introduction

T cells engineered with a chimeric antigen receptor (CAR-T therapy) is an adoptive cell therapy that recognize and targeted eliminate cancer cells expressing a specific target antigen. Since its first clinical application in Chronic Lymphoblastic Leukemia in 2010, CAR-T therapies have shown remarkable clinical efficacy, particularly against hematological malignancies, in numerous clinical trials^1,2^. However, CAR-T therapies are facing persistence issue due to several factors, such as reduced expression of the target antigen on cancer cells or dampened function of the transferred CAR-T cells. CAR-T cells, which are chronically stimulated by a cancer-specific antigen, will eventually differentiate into a dysfunctional exhausted state, characterized by reduced proliferative capacity and reduced effector function^3^.

High-throughput CRISPR−based genetic screens with either targeted or genome-wide library have become a widely−utilized method for identifying key regulators that can prevent CAR-T cell exhaustion and enhance their function^4^. Shifrut et. al performed a genome-wide loss-of-function screen (LOF) in primary human T cells and identified essential regulators for T cell receptor signaling and proliferation following stimulation^5^. Dong et, al performed the first *in-vivo* genome-scale CRISPR LOF screen of primary murine CD8^+^ T cells and identified regulators of CD8^+^ T cell tumor infiltration in a triple-negative breast cancer (TNBC) mouse model^6^. Schmidt et. al used both CRISPR activation (CRISPRa) and CRISPR interference (CRISPRi) genome-wide screens conducted in primary human T cells to identify the functional regulators of cytokine production in response to stimulation, which is often dysfunctional in autoimmune diseases and cancers^7^. Wang et. al conducted CRISPR LOF screens to directly identify essential regulators for cytotoxicity function in IL13Rα2-targeted CAR-T cells^8^. With the growing number of screens^9–14^ identifying genes that regulate specific T cell characteristics (e.g., survival, proliferation, or exhaustion) and functions (e.g., tumor infiltration, cytokine production, or cytotoxicity), the need for systematic databases that specifically map T cell gene functions has become increasingly urgent. Such resources are essential to advancing T cell-based therapies and addressing their rising demand.

In this study, we developed a large-scale, publicly accessible data repository called T cell perturbation genomics database (TCPGdb). TCPGdb will serve as a comprehensive resource depicting the functional effect of individual gene perturbation in T cells, covering all previously published T cell genome-wide screen datasets, as well as the transcriptional profiles of sorted T cells, CAR-T cells, T derived cancer cells and autoimmune disease datasets. In summary, TCPGdb provides a one-stop platform for searching, browsing, and visualizing T cell gene functions.

## Method

### CRISPR screen data collection and processing

T cell CRISPR pooled screen datasets were obtained from Gene Expression Omnibus (GEO)^15^, and previously published literature through searching Pubmed and Google scholar with keywords “CRISPR screen AND T cell”. To combine the CRISPR screen datasets, we retrieved the raw sgRNA data from all CRISPR screens and then processed each dataset using MAGeCK^16^ with uniform parameters. For datasets without raw data, the summarized data from original literature will be used. For those screens using mouse models, mouse genes were matched to their orthologous human genes using the biomaRt^17^ package in R. Detailed information on the included CRISPR screen datasets is provided in Table S1.

### RNA-seq and microarray data processing

For the RNA-seq data, raw sequencing reads were downloaded and unpacked using the SRA Toolkit (version 2.9.0-ubuntu64)^18^. Fastp was used to trim the adapter sequence and remove low-quality reads^19^. The trimmed reads were then mapped to the human reference genome GRCh38 (gencode.v45) and quantified at the gene level using the --quantMode GeneCounts option in STAR (v 2.7.10b)^20^. The count data were normalized to transcripts per million (TPM) in log2(x + 1) format.

The R packages affy or oligo were used to process the raw CEL microarray data. Robust multichip average (RMA)^21^ was applied to normalized microarray expression values. Probes were mapped to Ensembl IDs using the biomaRt package in R.

### Sorted T cells RNA-seq dataset

T cell bulk RNA sequencing data was obtained from two sources: the Database of Immune Cell Gene Expression, Epigenomics, and Expression Quantitative Trait Loci (DICE)^22^ and ABsolute Immune Signal deconvolution dataset (ABIS)^23^. DICE includes 967 samples covering 11 T cell subtypes, with gene expression data downloaded from DICE. The ABIS dataset comprises 46 samples including 12 T cell subtypes; the raw RNA-Seq data were downloaded and processed using standard RNA-Seq data processing methods as described above. T cell gene co-expression analysis was conducted using the DICE dataset.

### CAR-T bulk and single-cell RNA-seq dataset

We collected CAR-T datasets that include healthy/control samples and dysfunctional (exhausted, non-responding/impaired) samples. A total of five datasets were gathered: two bulk RNA-Seq datasets and three scRNA-Seq datasets from three different studies. Detailed information on these datasets is summarized in Table S2.

### T cell lymphoma/leukemia dataset

We obtained three T cell lymphoma/leukemia dataset featuring gene expression prognostic analysis from GEO: GSE19069 (n=147)^24^, GSE58445 (n=193)^25^ and GSE90597 (n=29)^26^. The array datasets were downloaded and processed using the microarray data processing procedures described later. Clinical and survival information were retrieved from the original studies or requested from the corresponding authors. Detailed information on the T cell lymphoma datasets is provided in Table S3.

### Transcriptomic profiling of autoimmune diseases

We obtained gene expression datasets profiling peripheral blood mononuclear cells for three types of autoimmune diseases from GEO: four multiple sclerosis (MS) datasets, three inflammatory bowel disease (IBD) datasets, and three systemic lupus erythematosus (SLE) datasets. The microarray and RNA-Seq datasets were downloaded and processed using the microarray data processing procedures described later. Detailed information on the collected autoimmune disorder datasets is summarized in Table S4.

### Gene set enrichment analysis

Gene set enrichment analysis (GSEA)^27^ was performed to evaluate the effect of CRISPR screen gene targets at the genepathway level. The analyses were conducted using the R package clusterProfiler^28^ with curated gene set of biological process(BP) category in gene ontology.

### Survival analysis

The survival analysis of a specified gene in T cell lymphoma/leukemia dataset was calculated using a Kaplan–Meier (KM) model through R survival packages. P values less than 0.05 were considered statistically significant.

### T cell perturbation score calculation

To overcome the bias in gene selection results from various T cell screening datasets, we introduced a T cell perturbation score (TPS score) to systematically measure the impact of specific genes on T cell function following CRISPR activation or interference. A gene is considered more impactful if it consistently ranks near the top across multiple datasets. We utilized the RRA algorithm^29^ to compute the significance score (ρ value) for each gene by integrating multiple MAGeCK results. To interpret both loss-of-function and gain-of-function screens in the same manner of enhancing T cell function, we integrated the MAGeCK results as follows: for scoring gene activation impact, we chose negative ranks towards function in knockout screens and positive ranks towards function in activation screens; conversely, for scoring gene knockout impact, we chose positive ranks towards function in knockout screens and negative ranks towards function in activation screens.

We first normalized the MAGeCK gene summary ranks into percentiles ranks U = (u_1_, u_2_, …, u_m_), where u_i_ = r_i_ /m (i = 1, 2, …, m), and m represents the total number of genes in the rank. Under the null hypothesis, where the percentiles follow a uniform distribution between 0 and 1, the k-th smallest value among u_1_, u_2_, …, u_m_ is an order statistic that follows a beta distribution. RRA computes the p-value for each gene based on the beta distribution in each rank. The significance score of the gene across ranks, the ρ value, is defined as ρ=min (p_1_, p_2_, …, p_n_), where n is the number of informative ranks. The TPS score for each gene was calculated with modified RRA ρ value as following:

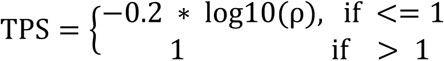

The TPS score ranges from 0 to 1. Genes with a TPS score near 1 are associated with a strong positive impact on T cell functions. Conversely, genes with a TPS score near 0 indicate minimal effect on enhancing T cell functions. The TPS score provides researchers with a quantifiable measure to evaluate the extent of a gene’s impact on T cell functions.

### LASSO model for CAR-T function prediction

LASSO, a widely used regularization method for handling multicollinearity and small datasets, was employed to build a regression model using CAR-T transcriptomic datasets. We utilized the Scikit-learn package in Python and applied Leave-One-Out Cross-Validation (LOOCV) to determine the optimal regularization parameter (λ).

### Database development

TCPGdb was powered by the Python Flask frameworks (https://flask-restful.readthedocs.io/). HTML, CSS were used for the rendering and interactive operations of the front-end pages. MongoDB Atlas (https://cloud.mongodb.com/) was used for storage of the processed data (https://www.mongodb.com/). The charts were manufactured by Echarts and matplotlib in python. Finally, the bioinformatics analyses were performed using R scripts. The R scripts usd are included as supplementary material. TCPGdb is hosted using Heroku and redirected to custom domain.

### Data availability

The data analyzed in this study were obtained from previously published papers. We have included references for each of the datasets used and all datasets are publicly available. The sources of all data sets were shown in Supplementary Tables.

## Results

### Data summary of T cell screen datasets used to generate TCPGdb

The TCPGdb database was generated from T cell CRISPR screen datasets in 13 published studies. These 13 studies included 8 CRISPR-activation screens, 2 open reading frame (ORF) gain-of-function (LOF) screens, 15 CRISPR-knockout screens, and 2 CRISPR-interference screens. Within the 27 included screens, 21 of them were genome-wide (**Fig. 1A**). The cell types targeted by the CRISPR screens included CD8+ T cells (n=12), CD4+ T cells (n=7), Treg cells (n=5), CAR-T cells (n=3), and mixed T cells (n=1). 12 of the 27 screens used mouse cells and the remainder (n=15) used human cells (**Fig. 1B**). The biological functions studied varied, including T cell killing, proliferation, survival, and the regulation of IL2, IFNG, and TNF expression. Most screens were conducted under *in vitro* conditions using genome-scale libraries (**Fig. 1B**).

**Figure 1.**
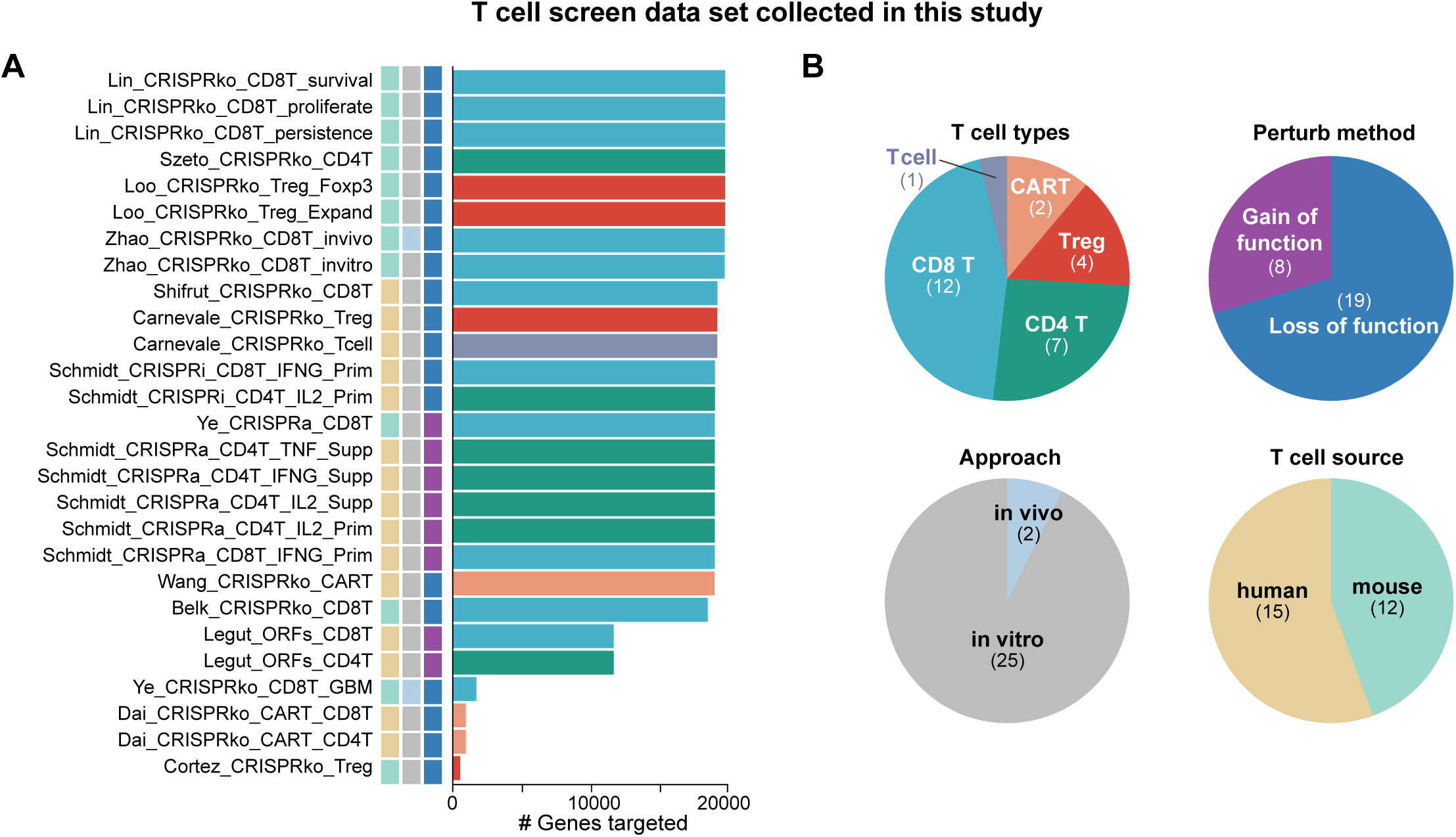
Summary of T cell screen datasets used to generate TCPGdb. (A) Bar plot of target gene library size in each screen experiment. (B) Pie chart showing the number of experiments belong to different subcategories.

### MAGeck re-analysis of CRISPR screen datasets used in TCPGdb

For 13 screens we were able to obtain the raw sgRNAs data. For these screens we re-analyzed the dataset using the MAGeCK pipeline with uniform parameters. Detailed process metrics can be found in Table S1. GSEA was used to evaluate the pathway enrichment based on gene-level results from MaGeCk, including targets identified through both positive selection and negative selection. Our re-analysis of all datasets revealed a large variability in the number of significant gene hits and pathways between screens, highlighting the impact of differences in experimental design, even among the same T cell types (**Fig. 2A**). Additionally, correlation analysis of the screening datasets revealed that among the 13 CD8+ T cell screen datasets, 15% of the comparisons between individual screens showed significantly positive correlations at the dataset level (12 out of 78 comparisons, p<0.05 and coefficients > 0.1). In contrast, only two significant negative correlations were identified (**Fig. S1A** and **Table S5**). For the CD8+ T cell proliferation screen datasets in particular we observed only positive correlation coefficients between screens (**Fig. 2B**), indicating the underlying similarities despite heterogeneity between the datasets. Furthermore, we observed shared targets hits between different screen datasets (**Fig. 2C**). For instance, among the six CD8^+^ T proliferation CRISPR knockout screen datasets, *ARIH2*, *CBLB*, *CD5*, *CD8A*, *CDKN1B*, *DGKZ*, *ELOB*, *MEF2D*, *RPRD1B*, *SOCS1*, *TMEM222*, *TNFAIP3* and *UBASH3A* were positively selected in Shrifut et al. and Carneveke et al., *LARP1* and *TP53* were both identified in Lin et al. and Zhao et al., *RC3H1* and *ZC3H12A* was enriched in the *in-vitro* and *in-vivo* datasets in Zhao et al. and *ARID1A* was identified in both Lin et al. and Carneveke et al., finally *RASA2* was found to enhance T cell proliferation in three datasets (Lin et al., Shifrut et al., Carnevale et al.) (**Fig. 2C**). We did not observe shared signaling pathways enrichment among the CD8+ T cell screen datasets.

**Figure 2.**
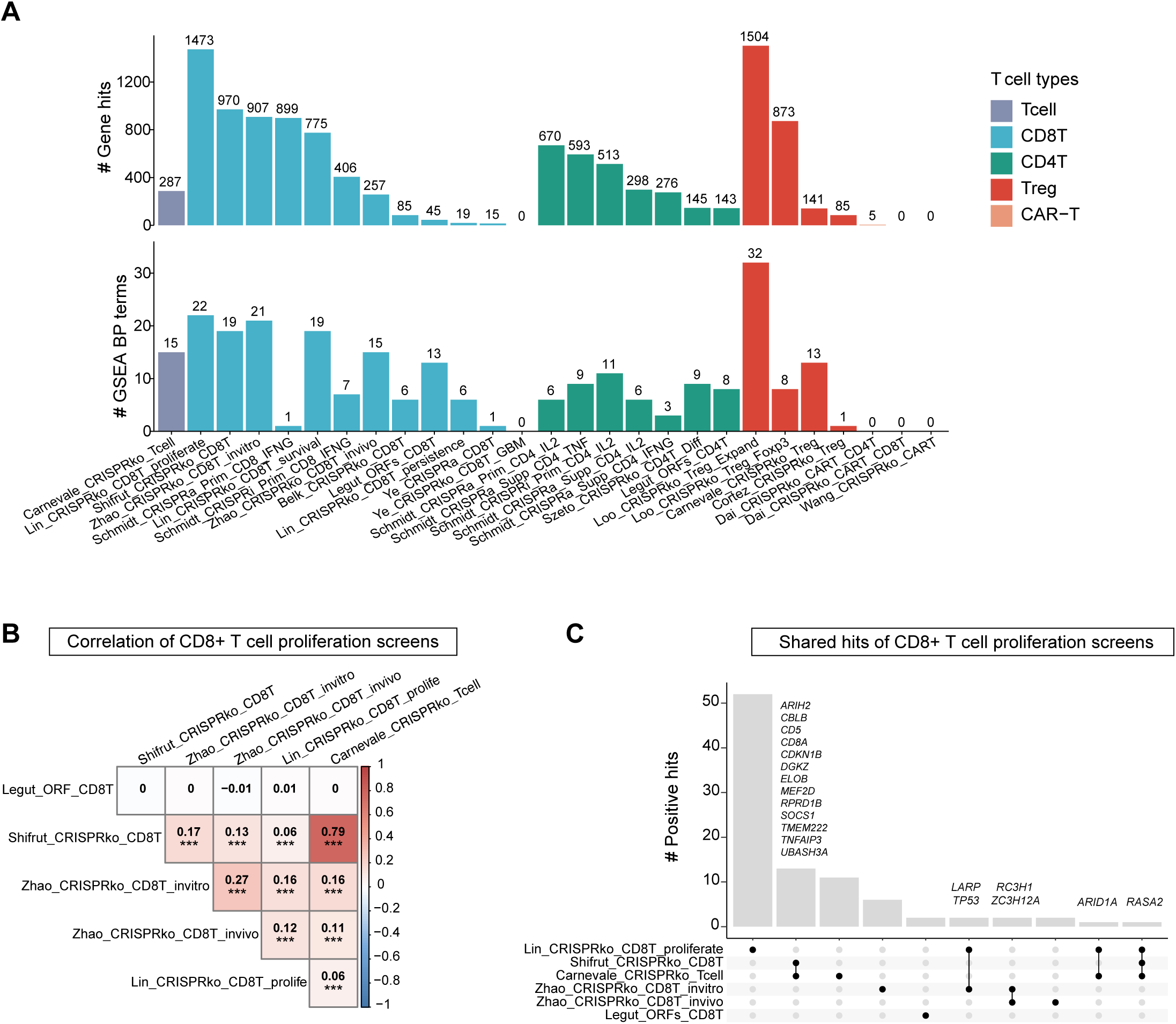
Summary of MAGeck analysis results for screen datasets used to generate TCPGdb. (A) Bar chart showing the number of differentially enriched genes (DEGs) (above), and GO terms enriched among identified genes (bottom) with significant statistical difference. Color of the bar indicates different T cell types. Level of statistical significance: DEGs (adjust pvalue <0.05); GSEA BP terms (pvalue < 0.05). (B) Correlation plot showing the correlaton of CD8+ T cell proliferation screen datasets by positive log fold change from MAGeCK analysis. The displayed values represent the Spearman correlation coefficient, with asterisks indicating the significance levels: * for p < 0.05, ** for p < 0.01, and *** for p < 0.001. (C) Upset plot showing the positive selected genes shared by CD8+ T cell proliferation screen datasets from MAGeCK analysis.

### Evaluation and characterization of genes with top T cell gene perturbation scores from activation and knockout CRISPR screens

After integrating the CRISPR screen datasets, we used a T cell perturbation score (TPS) metric to comprehensively characterize the effect of each gene perturbation on T cell function. The TPS score is calculated by summarizing multiple selection rank results obtained from MAGeCK, which allows a more intuitive understanding on the effect of each gene perturbation (both activation and inactivation) on augmenting T cell function (see Materials and Methods section for details). The calculated TPS score ranges from 0 to 1, with a higher TPS indicating that perturbing the gene has a greater impact on enhancing T cell functions, while lower TPS associates with less impact. We applied the TPS model to calculate the gene score under activation and knockout screen designs using CD8^+^ T cell, CD4^+^ T cell, Treg cell and CAR-T cell screening datasets. We observed positive correlations between the average of z-scored log fold change of genes and their TPS scores across each T cell types (**Fig. S1B**). We further analyzed the z-scored log fold change for the top 50 genes ranked by the TPS for both activation/gain-of-function (GOF) (**Fig. 3A**) and knockout/loss-of-function (LOF) (**Fig. 3B**) perturbation in CD8^+^ T cells. In addition to the overall impact on T cell function, we also used the TPS model to evaluate the gene potential in enhancing specific functions such as proliferation, cytotoxic killing, tumor infiltration, persistence, survival and interferon-gamma (IFNG) expression. The top 3 activation genes, *HSPA8*, *ALDOA* and *RPL5*, all showed a high TPS score for proliferation. *HSPA8* and *ALDOA* had a high TPS score in proliferation, persistence and survival, while *RPL5* also showed enhanced cytotoxic killing (**Fig. 3C**). For the top 3 knockout genes *TP53*, *CBLB* and *ZC3H12A*, inactivation of all three genes was associated with enhanced proliferation and persistence. Additionally, *TP53* was associated with survival, and *CBLB* associated with *IFNG* expression (**Fig. 3D**). Pathway enrichment analysis of the top 50 TPS activation genes showed that the top activation genes were mainly enriched in the RNA processing and translation–related pathways (**Fig. 3E**). In contrast, top knockout genes were mainly enriched in immune signaling and cell proliferation pathways, including immune response, T cell receptor signaling pathway, cell adhesion, intracellular signal transduction, regulation of cell cycle and negative regulation of cell proliferation (**Fig. 3E**). Similarly, we applied the TPS model on CD4^+^ T cell screening data and pathway enrichment analysis revealed that both top activation and knockout genes were enriched in T cell receptor signaling and immune response pathways (**Fig. S2A-C**).

**Figure 3.**
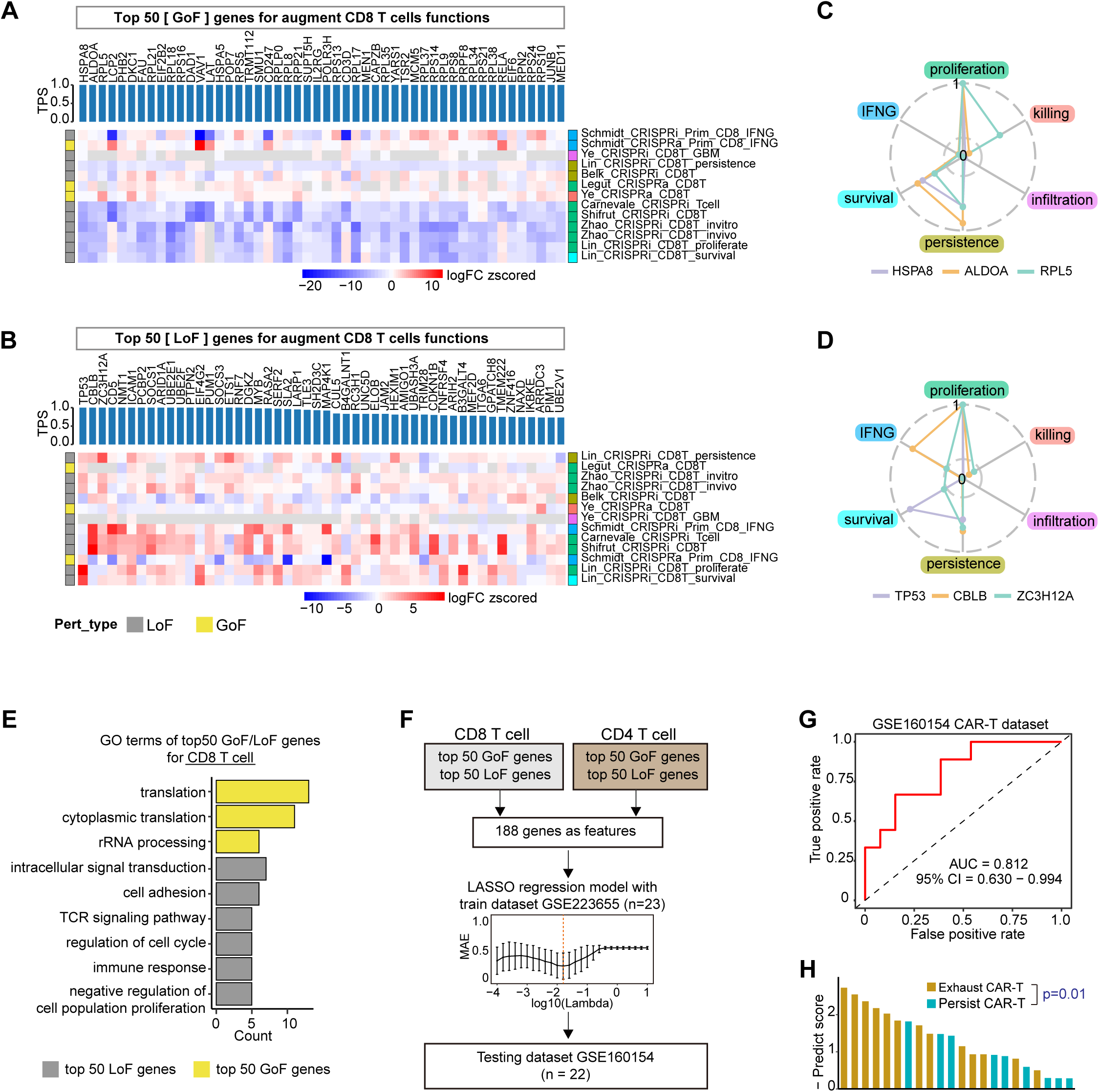
Summary of top genes and pathways ranked by T cell perturbation score (TPS) from 13 CD8+ T cell screen datasets. (A) Heatmap of z score fold change for top 50 gain-of-function (GoF) genes ranked by TPS from 13 CD8+ T cell screen datasets. (B) Heatmap of z score fold change for top 50 loss-of-function (LoF) genes ranked by TPS from 13 CD8+ T cell screen datasets. (C) Radar plots showing top 3 GoF genes and TPS score for gene potential in enhancing specific functional characteristics (proliferation, killing, infiltration, persistence, survival and *IFNG* expression). (D) Radar plots showing top 3 LoF genes and TPS score for gene potential in enhancing specific functional characteristics (proliferation, killing, infiltration, persistence, survival and *IFNG* expression). (E) GO terms enriched from the top 50 GoF and top 50 LoF in CD8^+^ T cells. Count represents the number of genes in the enriched terms. (F) Overview of the LASSO model training and testing procedure: A LASSO multivariate model was trained on T cell genes with top TPS scores to predict CAR-T putative function. The model was using leave-one-out validation to identify the optimized hyperparameters before applied to an independent CAR-T dataset (G) ROC curve analysis of LASSO model performance in prediction using the independent CAR-T dataset GSE160164. (H) Bar plot of prediction scores in the testing dataset, categorized by true labels. Statistical significance was assessed using the Wilcoxon rank-sum test.

We noticed that some genes showed a high TPS score in both CD8^+^ and CD4^+^ T cells. Among the top 50 TPS activation associated genes this included: *CD247*, *LAT*, *LCP2*, *RELA*, *RPS5* and *VAV1* (**Fig. S2D**). For the top 50 TPS inactivation associated genes this included: *CBLB*, *MAP4K1*, *SLA2* and *UBASH3A*. Furthermore, we tested the overlapping genes in two CAR-T bulk RNA-seq datasets, GSE160154 (CAR-T exhaustion) and GSE223655 (CAR-T response) and observed that the shared activation gene *RPS5* was significantly lower in exhausted CAR-T cells compared to controls, while the other five activation genes also showed a trend of decreased expression in exhausted CAR-T cells (**Fig. S2E**, upper panel). Additionally, *LAT* and *RELA* showed a statistically significant increase in CAR-T cells from responders compared to non-responders (**Fig. S2E**, bottom panel). Among the shared inactivation-associated genes, none were significant in exhausted CAR-T cells, but *CBLB* expression was increased in responder CAR-T cells (**Fig. S2F**).

We further developed multivariate models to predict CAR-T cell therapy outcomes, based on the top 50 GOF and LOF genes with the highest TPS scores from CD8^+^ and CD4^+^ T cell screens. Specifically, we trained a Least Absolute Shrinkage and Selection Operator (LASSO) logistic regression model using the GSE223655 dataset, which includes data from 12 responders and 11 non-responders to CAR-T therapy. We optimized the model parameters using leave-one-out cross-validation (Fig. 3F). We then tested the model’s performance on an independent dataset, GSE160154, which consist of 9 persistent and 13 exhausted CAR-T cell samples. The regression model demonstrated robust power in predicting CAR-T functionality with an AUC of 0.812 in the testing dataset (Fig. 3G). A Wilcoxon test revealed a significant difference in prediction scores between exhausted and persistent CAR-T cell samples (Fig. 3H). The successful validation of this TPS gene-based model underscores the robustness of these functional genes in influencing real CAR-T cell function, despite variations in engineered CAR designs.

### TCPGdb web interface and functional modules

The datasets and analyses described above were integrated into the TCPGdb database web interface. The website consists of three main functional components: T cell CRISPR screen view ("T-Screen"), T cell gene perturbation scoring ("TPS"), and T cell gene function explorer ("T-GEXP"). The CRISPR screen module includes summaries of gene hits from 27 T cell screen datasets and GSEA BP pathway term enrichments (**Fig. 4A**). The TPS module provides a scoring system for evaluating the impact of specific gene perturbations on both overall and specific T cell functions by integrating results from multiple CRISPR screen datasets (**Fig. 4B**). Notably, we also provide a radar chart indicating CD8 T cell functions in different aspects such as proliferation, survival, killing ability, infiltration, persistence, and expression of IFNG or IL2 (**Fig. 4B**).

**Figure 4.**
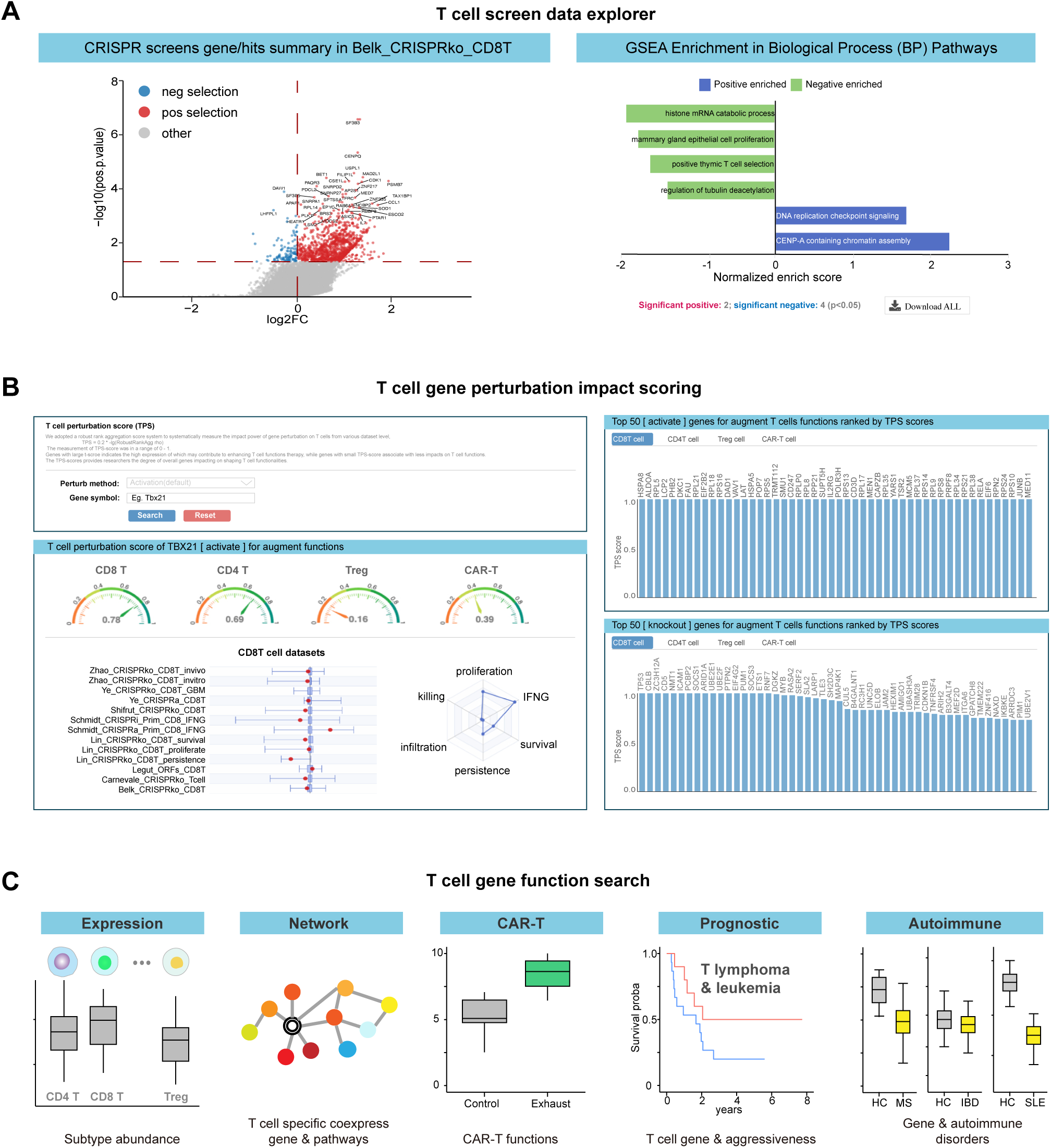
Overview of the TCPGdb web interface functional modules. (A) Gene summary and GSEA pathway analysis of T cell screen data included in this study. (B) Representative T cell perturbation score (TPS) page scoring system for an example gene. (C) TGExp search module in TCPGdb, include expression & co-expression, CAR-T performance association, T cell derived cancer prognosis and autoimmune diseases association.

To provide a comprehensive view of gene function specifically for T cells, we present the “T-GEXP” (T cell gene function explorer) module for exploring gene expression within different T cell subtypes, network analysis of co-expressed genes and pathways and links to T cell and CAR-T-related pathologies using existing clinical and experimental data (**Fig. 4C**). The query will return 5 main characteristic of the gene including: (1) Query gene expression: Expression profiles across 16 T cell subtypes using sorted T cell transcriptomic data from DICE^22^ and ABIS^23^. (2) T cell-specific co-expression (network): Co-expressed gene and pathway analysis using DICE T cell gene expression data (n=967). (3) Gene and CAR-T Cell functions: Analysis of gene impact on exhaustion and therapy responses using five CAR-T datasets: GSE160154, GSE223655, GSE151511, GSE160160, and GSE243325. (4) Gene and T cell cancer prognosis: Prognostic analysis for T cell-derived cancers using three T cell lymphoma datasets: GSE19069, GSE58445, and GSE90597. This reflects the potential of the query gene in enhancing T cell vigor and aggressiveness, inspired by Garcia et al., who discovered that naturally occurring T cell mutations can enhance engineered T cell functions^30^. (5) Gene and autoimmune Disorders: Analysis using transcriptomic profiles from peripheral blood mononuclear cells (PBMCs) in three autoimmune diseases (MS, IBD, and SLE) to explore associations between specific genes and autoimmune diseases, addressing potential side effects of T cell therapies. The T-GEXP search function is accessible via the T-GEXP search page, or the quick search box located at the top left corner of each page, making it convenient for users to retrieve information related to T cell therapies.

## Discussion

TCPGdb represents a pioneering and comprehensive resource designed for the analysis and visualization of T cell perturbation genomics within a single platform. We expect this resource will be beneficial for the scientific community to easily access and explore prior screen-based knowledge on the T cell function related gene targets. TCPGdb incorporates a wide array of data, including both gain-of-function and loss-of-function screens, conducted *in-vitro* and *in-vivo*, encompassing multiple T cell subtypes, engineered CAR-T screen datasets, and gene expression data from pure T cells and T lymphocyte-derived cancers. Through its database modules, researchers gain an informative gateway to explore the functional profiles of their target genes across various T cell perturbomics dataset collections.

While we have endeavored through TCPGdb to construct a comprehensive collection of T cell screen data encompassing various methodologies and phenotypes, we acknowledge the limitations arising from the cohort sizes of the datasets, which may introduce biases stemming from experimental conditions. Additionally, for the general function TPS-score, we lack a standardized approach to summarizing scores that effectively represent the overall impact of specific gene perturbations while integrating diverse T cell fitness phenotypes.

As the number of CRISPR screens conducted to investigate the effect of gene perturbation on T cell function continues to grow, we are also committed to continuously updating TCPGdb with new datasets. Our plan includes biannual updates to our database to incorporate all newly available datasets. Further datasets will increase the predictive accuracy of TCPGdb and potentially allow analysis of additional phenotypic characteristics beyond proliferation, survival, persistence, infiltration, killing and *IFNG* expression. Our web interface is publicly accessible and allows researchers with a limited bioinformatics background to easily search their gene of interest. Furthermore, we will introduce additional modules to facilitate seamless querying of T cell gene functionalities. TCPGdb stands poised as a valuable resource for studying T cell perturbation genomics and providing support to the rapidly evolving field of adoptive immune cell therapy.

## Supplemental Figure Legends

**Figure S1.**
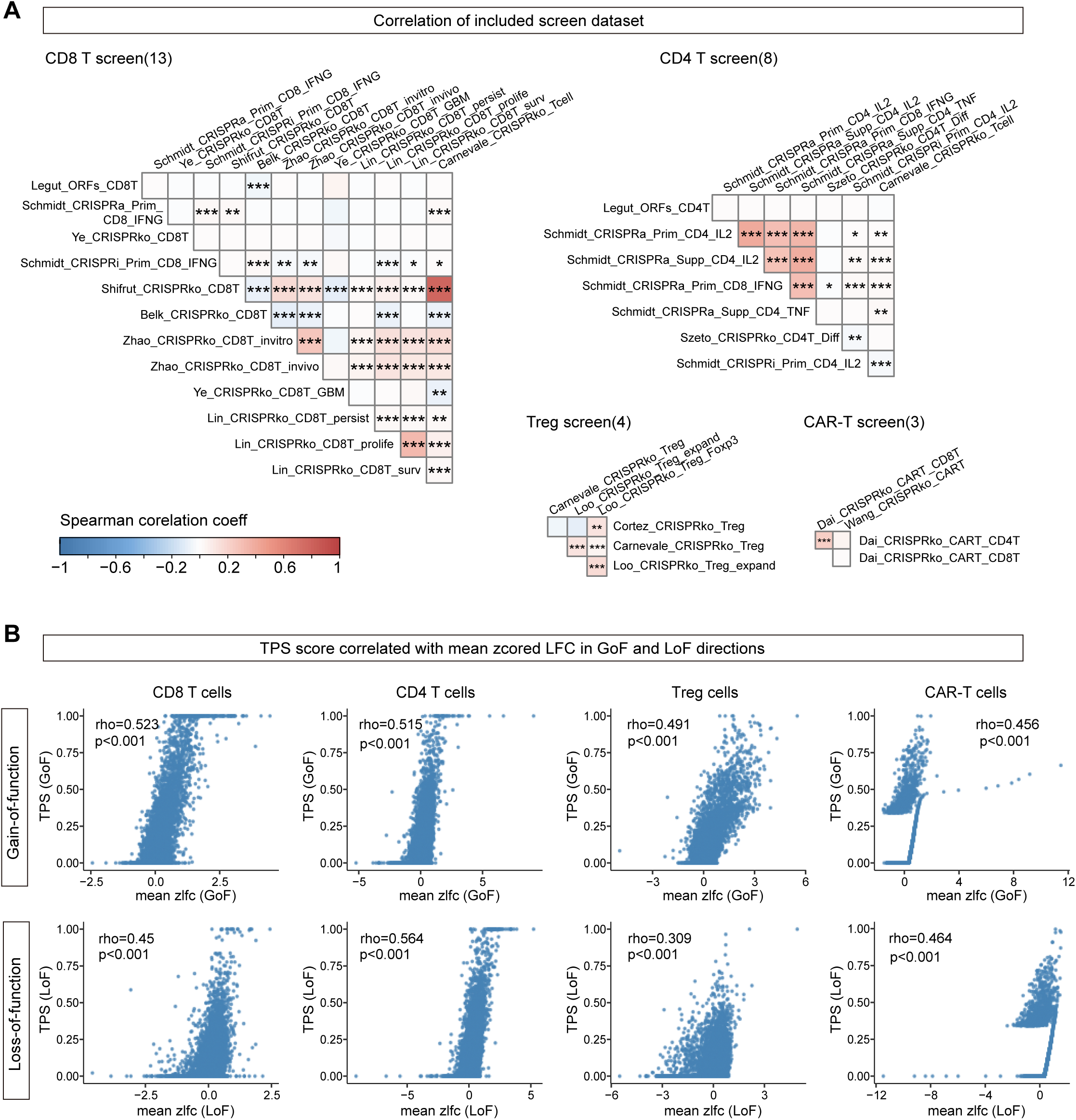
Correlation of T cell screen data sets and TPS scores. (A) Overview of screen data correlation in CD8+ T cells, CD4+ T cells, Treg cells, and CAR-T cells. To incorporate both loss-of-function and gain-of-function screens in a consistent manner, we integrated the MAGeCK results as follows: for knockout and interference screens, we chose negative logFC values towards augmenting function; conversely, for activation and open reading frame (ORF) screens, we chose positive logFC values towards augmenting function. Correlation between two experiments was assessed using Spearman correlation. Specifically, for the Wang_CRISPRko_CART dataset lacking raw data and logFC values, we used the enrichment score as an alternative. The displayed values represent the Spearman correlation coefficient, with asterisks indicating the significance levels: * for p < 0.05, ** for p < 0.01, and *** for p < 0.001. (B) Correlation of TPS score and z-scored log fold change in four T cell types. The correlation between TPS scores and mean z-scored logFC was measured using Spearman correlations. Mean z-scored logFC was calculated using the direction of logFC consistent with the ranks used in calculating TPS scores.

**Figure S2.**
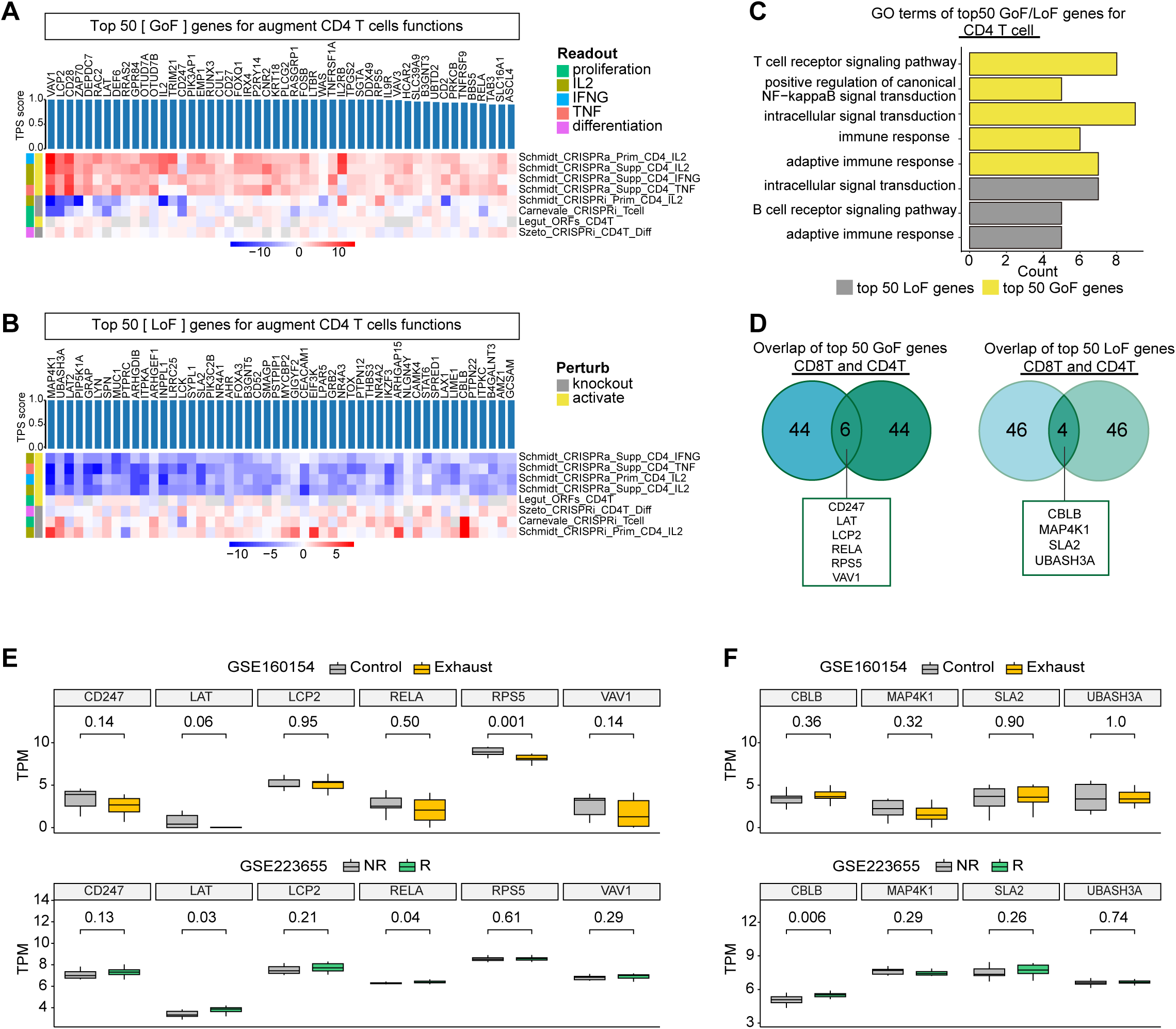
Summary of top genes and pathways ranked by T cell perturbation score (TPS) from CD4+ T cell screen data. (A) Heatmap of z score fold change for top 50 gain-of-function (GoF) genes ranked by TPS from 8 CD4+ T cell screen datasets. (B) Heatmap of z score fold change for top 50 loss-of-function (LoF) genes ranked by TPS from 8 CD4+ T cell screen datasets. (C) GO terms enriched from the top 50 GoF and top 50 LoF in CD4^+^ T cells. Count represents the number of genes in the enriched terms. (D) Venn diagram showing the overlap of top 50 GoF gene from CD8^+^ T cell and CD4^+^ T cells (G) screen data.| (E) Boxplot of overlapped GoF genes (showed in Fig. 3F) in CAR-T transcriptomic dataset. The upper panel shows *CD247*, *LAT*, *LCP2*, *RELA*, *RPS5* and *VAV1* expression in GSE160154, the CAR-T exhaustion datasets; the bottom panel shows expression of the six genes in GSE223655, the CAR-T response performance data. (F) Boxplot of overlapped LoF genes (showed in Fig. 3G) in CAR-T transcriptomic dataset. The upper panel shows *CBLB*, *MAP4K1*, *SLA2* and *UBASH3A* expression in GSE160154, the CAR-T exhaustion datasets; the bottom panel shows the four gene expression in GSE223655, the CAR-T response performance data.

## Table legend

**Table S1. T cell screen data set include in this study.**

**Table S2. CAR-T therapy data set include in this study.**

**Table S3. T-cell lymphoma/leukemia data set include in this study.**

**Table S4. Autoimmune disorders data set include in this study.**

**Table S5. Correlation of 13 CD8+ T cell screen datasets.**

## Notes

### Competing Interest Statement

The authors have declared no competing interest.

